# Aminoglycoside antibiotics perturb physiologically important microRNA contributing to drug toxicity

**DOI:** 10.1101/137935

**Authors:** Gopal Gunanathan Jayaraj, Soundhar Ramasamy, Debojit Bose, Hemant Suryawanshi, Mukesh Lalwani, Sridhar Sivasubbu, Souvik Maiti

## Abstract

miRNAs are key non-protein coding regulators of gene expression in various pathophysiological conditions. Targeting miRNA with small molecules offer an unconventional approach, where clinically active compounds with RNA binding activity can be tested for their ability to modulate miRNA levels and thus for drug repositioning. Aminoglycoside antibiotics are highly effective microbicidal RNA binding molecules that bind to prokaryotic rRNA secondary structures. Here, we report that specific subsets of miRNA can be modulated by aminoglycosides. However, ototoxicity (cochlear and vestibular) and nephrotoxicity of multiple origins resulting from prolonged use are a well-known disadvantage of aminoglycosides. Mature non-coding RNAs and their precursors can present off-target sites, by forming secondary structures that resemble ribosomal RNA, thus providing an additional molecular basis for the toxicity of aminoglycosides. Using in vitro, in cellulae and physiological responses, we provide evidence for the direct functional perturbation of the miR- 96 cluster leading to selective cell death in neuromasts- the zebrafish equivalent of cochlear hair cells, by Streptomycin, a prototype aminoglycoside antibiotic, thus contributing to the observed ototoxicity. Our observations, collectively underscore the importance of re-evaluating RNA binding drugs for their off-targeting effects in the context of miRNA and other functional non-coding RNA.

## INTRODUCTION

The knowledge that drugs prescribed as medicines, can cause adverse side effects due to various reasons is not new and has been explored in past extensively. Avoiding drug induced side effects/toxicities has emerged as the prime criteria in careful and meticulous evaluation of new drugs which are designed and tested for therapeutic interventions. Although most mechanisms of drug induced toxicities have been explained and categorized on the basis of the drug-target relationships, several caveats exist which prevents a comprehensive understanding of these adverse side effects. Drugs targeting proteins have been explored to a great extent and their side effects well studied. This, however, is not true for drugs which target RNA. Historically, the drugability of RNA has been demonstrated using well studied aminoglycoside antibiotics which have been an integral class of antibacterial agents for the past half a century. However, knowledge about aminoglycoside drug-target RNA interactions has been limited owing to their apparent specificity to prokaryotic systems which now have been questioned due to their adverse side effects(Begg and Barclay, 1995). It is also evident that most of the RNA binding, candidate drugs were tested for their efficacy and their side effects recorded without the knowledge of the full spectrum of the transcriptome which was earlier thought to only consist of mRNA, tRNA and rRNA. Extensive technological advances like next generation sequencing have uncovered the functionality of the human genome with unprecedented resolution, where now it is known that most of it is transcribed as functional non-coding RNA(Jacquier, 2009). microRNAs are one such class which serve to function as modulators of mammalian gene expression and have been extensively characterized as an indispensable layer of cellular regulatory molecules(Bartel, 2009). miRNAs form a complex network of gene regulators with a single miRNA estimated to control hundreds of mRNAs(Krol, et al., 2010). Their causal roles in disease pathology are well understood(Mendell and Olson, 2012) leading to the understanding that expression imbalance of miRNA are associated with a perturbation of cellular homeostasis. miRNA biogenesis involves the processing of longer precursor molecules which are enrich in secondary structures. Conventional methods of miRNA knockdown using chemically modified antisense-oligonucleotides have been explored extensively but face the challenges of modes of delivery, biostability and biodistribution. This calls for the development of more alternative and non-conventional methods to target miRNA. Small molecules targeting RNA chemical and structural space provide one such timely opportunity (Jayaraj, et al., 2015). We here explore the potential of well characterized RNA binding drugs (using aminoglycoside antibiotics as an example) to cross react with miRNA and characterize in detail the effects of Streptomycin which we previously showed to be a potential inhibitor of miR-21 function (Bose, et al., 2012). Streptomycin functions as an effective antibiotic via binding to the stem-loop structure (Hong, et al., 2014; Leclerc, et al., 1991) of the 16S rRNA of prokaryotic 30S ribosome resulting in inhibition of protein synthesis and mistranslation (Kohanski, et al., 2008), ultimately leading to the death of the microbial cells. Classically Streptomycin has been used to treat tuberculosis, although its widespread usage is limited due to its associated side effect, hearing loss or ototoxicity and to a lesser extent nephrotoxicity (Begg and Barclay, 1995). Ototoxicity of Streptomycin (and aminoglycosides in general) has been explored over four decades and ascribed many different mechanisms of which disruption of mitochondrial ribosome function (owing to the similarities to prokaryotic ribosome) has been explored in detail (Hobbie, et al., 2008; Prezant, et al., 1993). Other studies indicate that aminoglycosides can disrupt ER homeostasis by direct binding to proteins (Karasawa, et al., 2010). They also cause ototoxicity by inhibiting chaperoning activity of HSP73 and Calreticulin (Horibe, et al., 2004; Miyazaki, et al., 2004). However subsequent studies showed that calreticulin binding is rather protective than toxic (Karasawa, et al., 2011). Aminoglycosides have also been shown to inhibit cytosolic ribosomal activity and protein synthesis (Francis, et al., 2013), whereas attempts to reduce mitochondrial binding yielded designer aminoglycosides which selectively inhibit cytoplasmic protein synthesis contradicting the previous finding (Shulman, et al., 2014). A more detailed review of the various intracellular mechanisms of aminoglycoside toxicity reported so far has been discussed elsewhere (Karasawa and Steyger, 2011). Since the primary target of aminoglycosides are stem-loop structures, we hypothesized that aminoglycosides can bind to and disrupt function of physiologically relevant miRNA.

MicroRNA were recently shown to be important for the development of hearing, when the evolutionarily conserved role of miR-96 cluster was delineated (Mencia, et al., 2009). It was observed that loss of miR-96 function leads to progressive hearing loss (Lewis, et al., 2009; Mencia, et al., 2009). More recently, thorough surveys revealed expression of several other miRNA in the mammalian inner ear (Rudnicki and Avraham, 2012; Ushakov, et al., 2013). However it remains that the function of miR-96 cluster is evolutionarily conserved prompting our examination of this cluster in particular. We show that Streptomycin induced ototoxicity could be a result of perturbation of miRNA-96 cluster function (amongst other miRNA) amongst other known factors. Zebrafish has been established and used a model organism to screen for potential chemical agents which can protect against drug induced hearing loss (Bowman and Zon, 2010). This is due to the fact that the lateral lining of the developing zebrafish show extensive functional and molecular similarities (Whitfield, 2002) to the mammalian inner organs. Using a series of cellular transcriptional responses complemented by biophysical and biochemical validation, we uncover a hitherto unknown mode of aminoglycoside action by global perturbation of microRNA levels which may result in new indications for aminoglycosides. We extend our analyses to an in vivo scenario using zebrafish as a model system and explain alternate modes of Streptomycin induced ototoxicity as an example.

## RESULTS

### Aminoglycosides induce differential expression of miRNA

To systematically deconvolute the effects of aminoglycosides, we initially used a prototypic aminoglycoside, Streptomycin, to screen for global effects of miRNA perturbation. When such small molecules are tested, with a view of inhibiting miRNA function, the question of specificity always takes prime importance. To address this, we performed a cell based assay, where we quantified levels of 380 miRNA (of which are at least 100 are present in functional abundance in MCF-7). We isolated total RNA from cells treated with Streptomycin at a concentration of 10 μM for 24 hours. Relative RT-PCR quantification of miRNA levels revealed that many miRNA showed differential expression. We observed a non-uniform distribution of fold change (for top 50 miRNA expressed) indicating miRNA specific responses upon treatment with Streptomycin. (See Fig S1a). Although we observe that Streptomycin downregulates a number of miRNA (of which miR-21 is significantly downregulated), upon applying an arbitrary cutoff of 2 fold (log_2_ transformed) we obtain 13 miRNA (hereafter referred to as hits). We repeated these analyses using a panel of 5 other related (natural and semisynthetic) aminoglycosides- Dihydrostreptomycin, Gentamycin, Neomycin, Amikacin and Tobramycin. All the drugs were treated at the same dosage (10 μM) to MCF7- cells and the miRnome wide transcriptional response recorded (See Fig S1b). For the ease of analysis, we restricted our analysis to functionally relevant miRNA (See Fig S2 and Supplementary note 1) and obtained an expression atlas across 6 drugs (Fig 1a). We observed that for most miRNA, there is a differential response depending on the aminoglycoside drug administered. Correspondingly, each drug generated a global but differential response across the functional miRnome (Fig 1b). We then filtered the dataset to obtain hits above the 2 fold cutoff for each small molecule drug treatment. We then constructed a network of small molecule drug and miRNA transcriptional responses, based on the lists of downregulated miRNA generated (Fig 1b and 1c). From the network properties we note that, Streptomycin, Gentamycin, and Dihydrostreptomycin have the least degrees (i.e. connected nodes). Streptomycin, Gentamycin, and Dihydrostreptomycin cause fewer global effects (Fig 1a and 1b), suggestive of a better starting point as a scaffold for modification to improve specificity and selectivity. Further these molecules may serve as valid predecessors of potential small molecule modulators of miRNA function.

**Figure 1.**
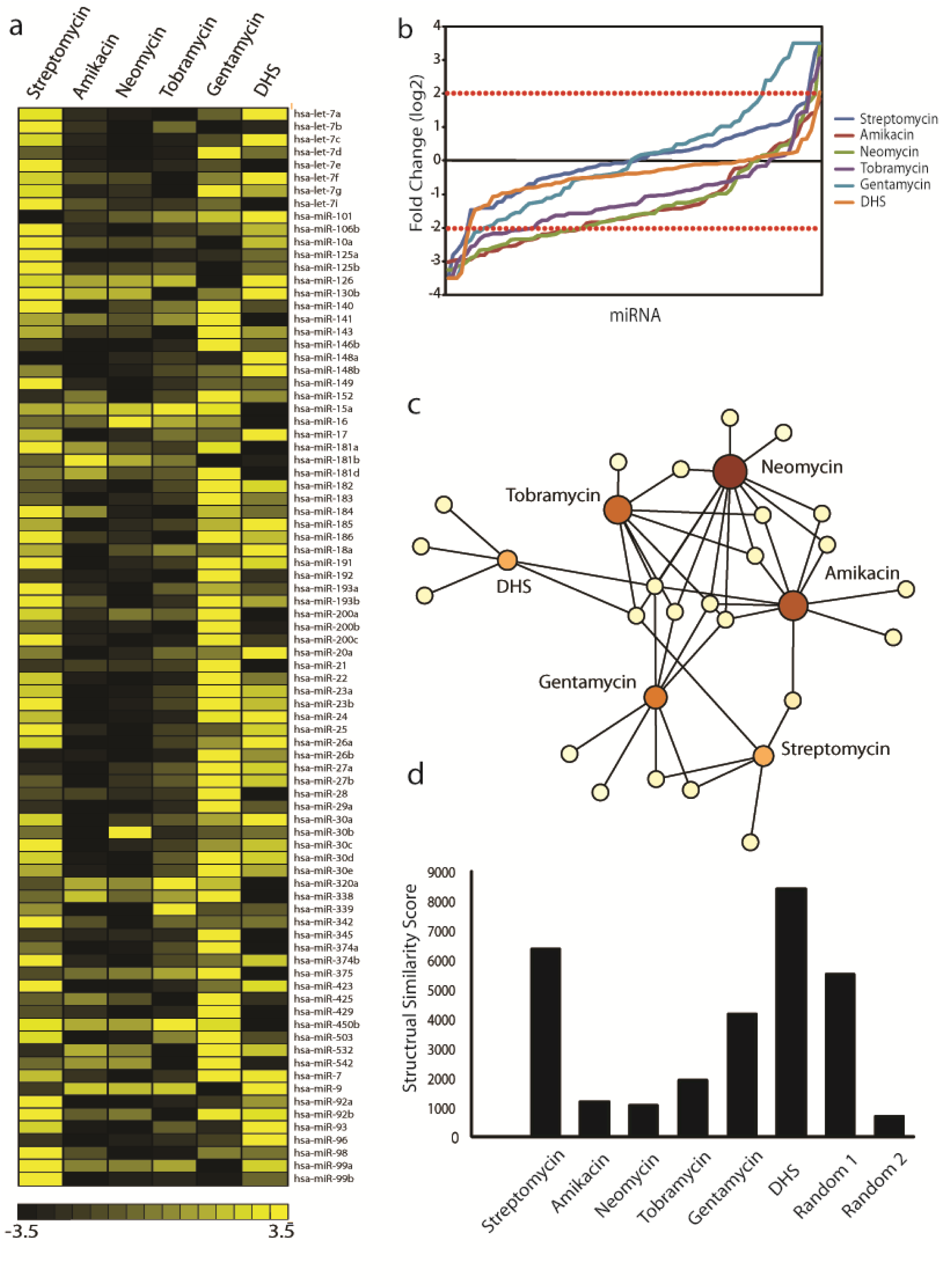
a) Expression changes of the functional 85 miRNA expressed in MCF7 in response to Streptomycin (10 μM, 24h). All the values represented are average of 2 biological replicates corrected for error (see Methods). b) Global distribution plot showing effect of each aminoglycoside drug. The dotted lines indicate the 2 fold cut-off applied to identify the hits. c) The complete MCF7 miRNA response network of all six small molecule aminoglycoside drugs. The coloured circular nodes represent the aminoglycosides, while the empty circular nodes represent miRNA. The number of edges connecting the coloured nodes form the number miRNA shared or unique and indicate the degree of each node. Dihydrostreptomycin, Streptomycin and Gentamycin have the least degree, making them suitable starting scaffolds. d) Plot to show the structural similarity score computed by the LocaRNA server. Higher the score (y-axis), indicates a more conserved structure between the miRNA (Random set 1 = 5 miRNA, Random set 2 = 31 miRNA - minimum and maximum number of miRNA hits obtained). In all cases, DHS=Dihydrostreptomycin.

It is usually impossible to distinguish the direct and indirect effects of miRNA expression changes upon small molecule treatment, however we rationalized that these hits may harbor similar secondary structure attributes. Therefore, we analyzed the local structural similarity(Will, et al., 2012) of the precursor miRNA between these hits for all the drugs and gauged that indeed these miRNA have elements of structural similarity as compared to randomly selected miRNA (Fig 1d, Fig S3a-S3f). We gauged that there is a partial dependence on structure and a more thorough investigation would be essential to separate out the observed effects.

These results led to two major implications, Firstly, that there are specific miRNA subsets in the miRnome which can respond to specific aminoglycoside treatment and it is imperative that these small molecules are to be further scrutinized to increase specificity and selectivity. Secondly, the clinical aminoglycoside antibiotic administration could modify the expressed miRnome thereby contributing to their associated toxicities. We also noted that almost all the drugs cause an increase in expression in a subset of miRNA when compared with the untreated cells. Notably for Streptomycin alone, some of the miRNA (Fig S1a) are up regulated (24 hits, 2 fold cutoff, see Supplementary Table S1) which may be due to the differential effects of the Streptomycin binding and increased dicing efficiency (See supplementary note 2) among other reasons.

### miRNA dysregulation is not due to a generic perturbation of miRNA biogenesis pathway

It is know that aminoglycoside administration induces mitochondrial stress, disruption of ER homeostasis amongst other effects on mammalian cells (Kalghatgi, et al., 2013; Karasawa and Steyger, 2011; Oishi, et al., 2015). Such a stressed state is typically known to have a very strong effect on miRNA regulation (Emde and Hornstein, 2014; Leung and Sharp, 2010). We sought to understand if the observed effect is due to a general failure of the miRNA biogenesis machinery. To this end, we treated MCF7 cells at 2X and 20X (where X=10μM) streptomycin and measured the levels of key miRNA pathway genes Dicer, AGO1, AGO2, AGO3 by quantitative RT-PCR. We observed no significant changes in expression levels at 2X concentration (Fig 2a). This held true even at 20X concentration of Streptomycin. Independently we examined a recently published data (Oishi, et al., 2015), where HEK293 cells were treated with Geneticin (a derivative of Gentamicin) and measured the response using microarrays. Meta-analysis of the data revealed that most of the miRNA biogenesis pathway genes were not significantly perturbed (Fig 2b). Taken together, miRNA dysregulation appears to be independent of any perturbation of a global factor governing miRNA biogenesis.

**Figure 2.**
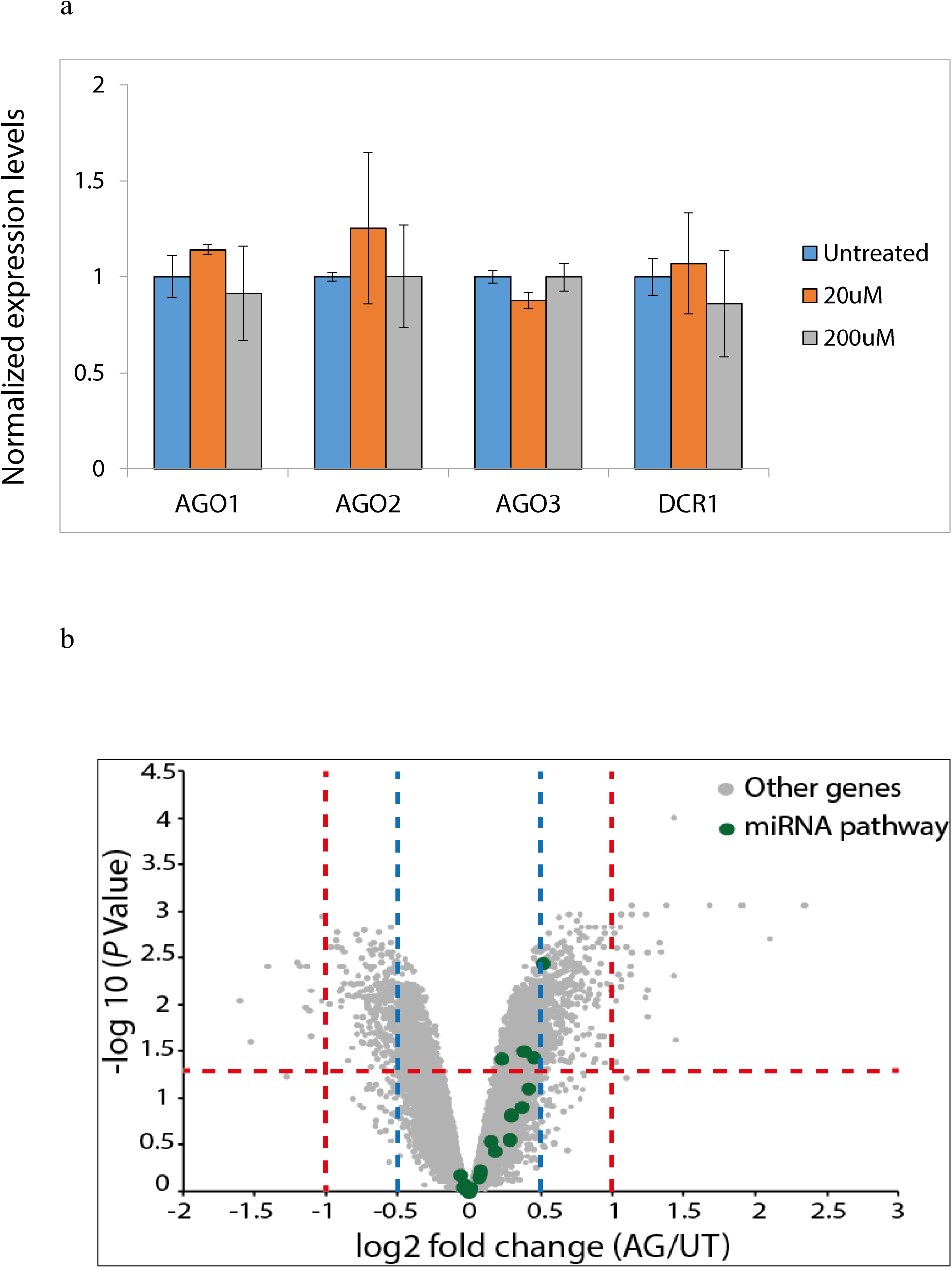
a) qRT-PCR showing the levels of key miRNA biogenesis pathway genes in response to Streptomycin treatment. Even at 200μM concentration, there seems to be no significant change in their levels. b) Volcano plot of data from GSE57198 showing the global gene expression profile of HEK293 cells in response to Geneticin (a derivative of Gentamicin). Most of the miRNA biogenesis pathways genes are not significantly altered.

### Functional studies hint at miRNA perturbation as a potent contributor to side-effects of aminoglycosides

To systematically analyze the effects of each drug molecule, we prepared response sets for each treatment with a reduced stringency to capture more biologically relevant information. We started with Streptomycin, and a detailed analysis of function revealed an unexpected possibility of miRNA perturbation related to the side-effects of the drug. We observed the two miRNAs - miR-96 and miR-182 are downregulated by approximately 1.5 fold. Interestingly, these miRNA belong to a cluster (miR-96, miR-182 and miR-183) whose functions are indispensable for the development and maintenance of hearing function(Rudnicki and Avraham, 2012). Other studies in mice implied that loss of function of miR-96 and cluster members resulted in abnormal development of the inner ear leading to hearing loss (Lewis, et al., 2009). miRNA are known to have multiple roles based on the tissue of expression and functionally diverse repertoire of their target mRNA(Sood, et al., 2006). miR-96 is also known to function by promoting cellular proliferation and is expressed significantly enough in MCF7 (Guttilla and White, 2009). We rationalized that, if Streptomycin lowers expression of miR-96 and other relevant miRNA in the cellular model, it is possible that they may be downregulated in the case of systemic administration of Streptomycin, due to its continued usage for the treatment of Tuberculosis despite its extensively documented ototoxicity leading to hearing loss (Mitnick, et al., 2008). Although, previous reports converge on the mitochondrial damage - induced cell death as the major contributor (among other possibilities) to hearing loss, based on the observations, we asked if the clinical manifestation of drug’s side-effect can also funnel via functional perturbation of miR-96 cluster.

### miR-96 cluster members are perturbed in a whole animal physiological model by Streptomycin administration

We sought to test if miR-96 downregulation occurred in a physiological model. We chose zebrafish as a model system to test our hypothesis. As mentioned before zebrafish lateral line neuromasts share extensive similarity in function and structure to the mammalian inner ear and have been used for chemical screens for compounds which can protect against ototoxic drugs (Owens, et al., 2008). Additionally, the roles of miR-96 and other cluster members have been shown to regulate the levels of hair cells in developing zebrafish and regulate sensorineural fates (Li, et al., 2010) in a manner similar to mammalian systems. Knockdown of these miRNA led to loss of hair cells and other phenotypic abnormalities associated with underdeveloped hearing capacity (Li, et al., 2010). We also noted that miR-96, miR-182 and miR-183 were conserved across species (Rudnicki and Avraham, 2012). A conservation plot based on the precursor miRNA (Fig. 3a), shows the extensive similarity of the cluster members.

**Figure 3.**
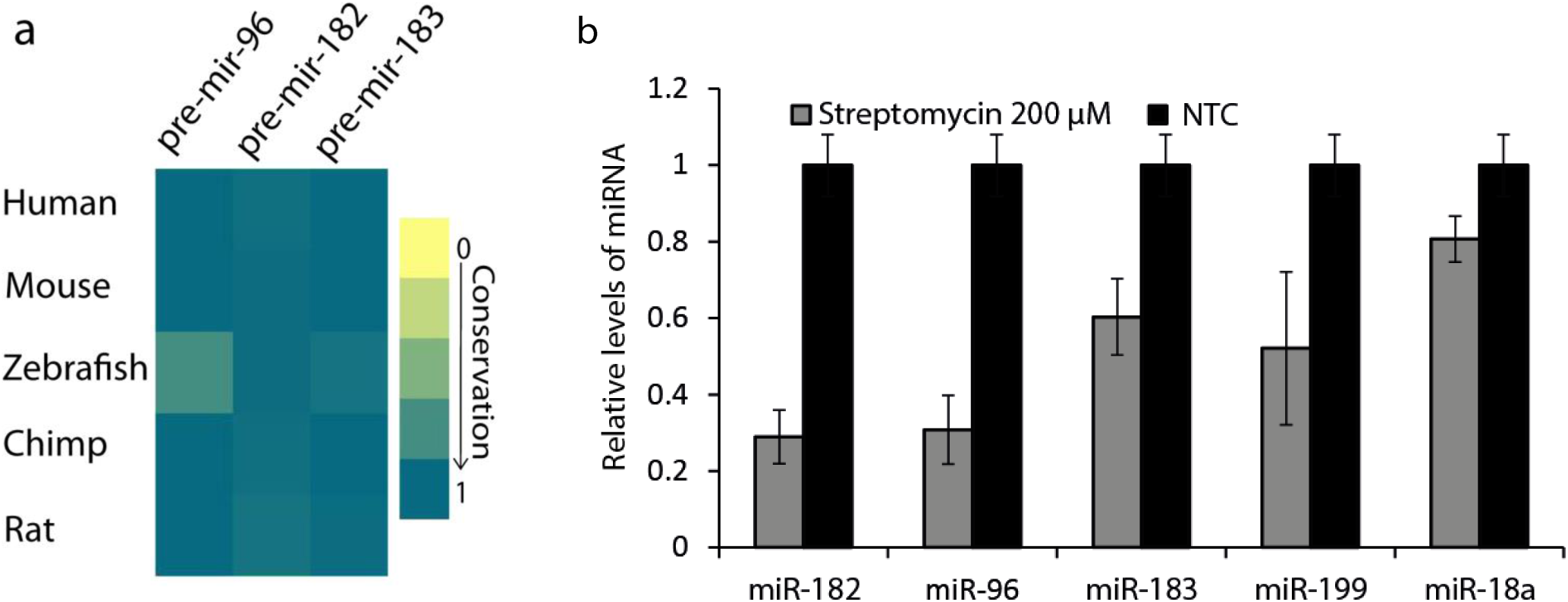
a) Conservation plot to show sequence and structural conservation of miR-96 cluster members across species. ZebrafishmiRNA shows a 75% structural and sequence similarity to human miRNA making it amenable to experimentation. b) qRT-PCR (average of 2 replicates from total profile) showing the decrease in levels of the miR-96 cluster members (and other conserved hearing miRNA reported in literature) in zebrafish embryos treated with Streptomycin.

To begin with, we assayed for the loss of hair cells on aminoglycoside treatment to confirm the viability of the model system. The hair cells of the neuromasts can be easily stained with vital dyes like DASPEI (a mitochondrial potentiometric dye) and thus visualized making the system amenable for testing the effects of aminoglycosides on the viability of the hair cells as previously described (Owens, et al., 2008). We observed a substantial loss of signal from the lateral line neuromast hair cells in 5dpf embryos treated with 200μM Streptomycin for 1 hour compared to the non-treated control set. This provided a starting platform to derive phenotypic readouts to compare miR-96 cluster knockdown and aminoglycoside treatment. To assess if the cluster miRNA levels are perturbed due to Streptomycin treatment, we quantified their levels through q-RT-PCR. We noted that miR-96 cluster members and other miRNA reported to be associated with hair cell development, were appreciably downregulated (Fig 3b). We then quantified global miRnome changes using a customized miRnome (248 miRNA) readout assay (Methods). Concordant to our results obtained in the cellular model, we observed a very asymmetric change in expression levels with many miRNA being downregulated while others being upregulated (Fig S4).

### Streptomycin binds directly to pre-miR-96 thereby reducing its dicing efficiency

To establish that aminoglycoside induced ototoxicity could channel via miR-96 cluster perturbation, we performed a series of in-vitro experiments. First, we addressed the question of whether Streptomycin actually binds to any of the members of the miR-96 cluster. To this end we performed S1 nuclease footprinting to determine if Streptomycin and other aminoglycosides confer any protection. Streptomycin showed protection for miR-96 cluster (Fig S6abc) whereas other aminoglycosides show no or negligible protection to this cluster (data not shown). Presence of Streptomycin in increasing concentrations reduced the cleavage of the RNA by S1 nuclease (Fig S6abc) indicating the binding of Streptomycin close to the stem loop junction (protected residues are highlighted in red in Fig S6). We note that this protection was most substantial for miR-183, followed by miR-96 and miR-182 suggesting that all 3 miRNAs are affected to varying degrees.

Detailed investigation of the binding affinity and kinetic parameters were done using Surface Plasmon Resonance (SPR) experiments. The binding affinities of pre-miR-96 and pre-miR- 182 (human and zebrafish) to Streptomycin were comparable and were in the ranges of values reported in literature for aminoglycoside binding (Fig S5 and Supplementary Table S2). In particular the binding affinity of Streptomycin to human pre-miR-96 was 6.50E+07 M^-1^.

Having established that Streptomycin can bind to pre-miR-96, we probed for its potential to perturb the maturation process by blocking the dicing reaction. We observed that for the aminoglycosides and miRNA tested, Streptomycin was able to prevent dicing to the maximum extent (Fig S6d).

### Streptomycin treatment produces a phenocopy of miRNA knockdown

Next we asked if the effects of Streptomycin treatment are similar to those of miRNA knockdown. To this end, we phenotypically profiled zebrafish embryos upon treatment with different aminoglycosides including Streptomycin. As mentioned before, the hair cells of the lateral line neuromasts can be visualized with DASPEI. We treated 5 dpf embryos with Streptomycin and other aminoglycosides at a concentration of 200μM. Fluorescence microscopy revealed a uniform loss of signal from the hair cells concentrated at the neuromasts (Fig 4a). For a more clear visualization of the hair cell bundles, we imaged embryos using confocal microscopy (Fig 4a inset). The treatment of each of the aminoglycosides except for Ampicillin (a non-aminoglycoside antibiotic) caused a considerable loss of signal (Fig S7a). We investigated if there is a corresponding decrease in miRNA levels. In situ hybridization using probes against miR-96, miR-182 and miR-183 showed a complete loss in signal in embryos treated with Streptomycin (Fig 4a inset and Fig S7b). Collectively, these results suggest a functional connection between loss of hair cells and reduction in miRNA levels.

**Figure 4.**
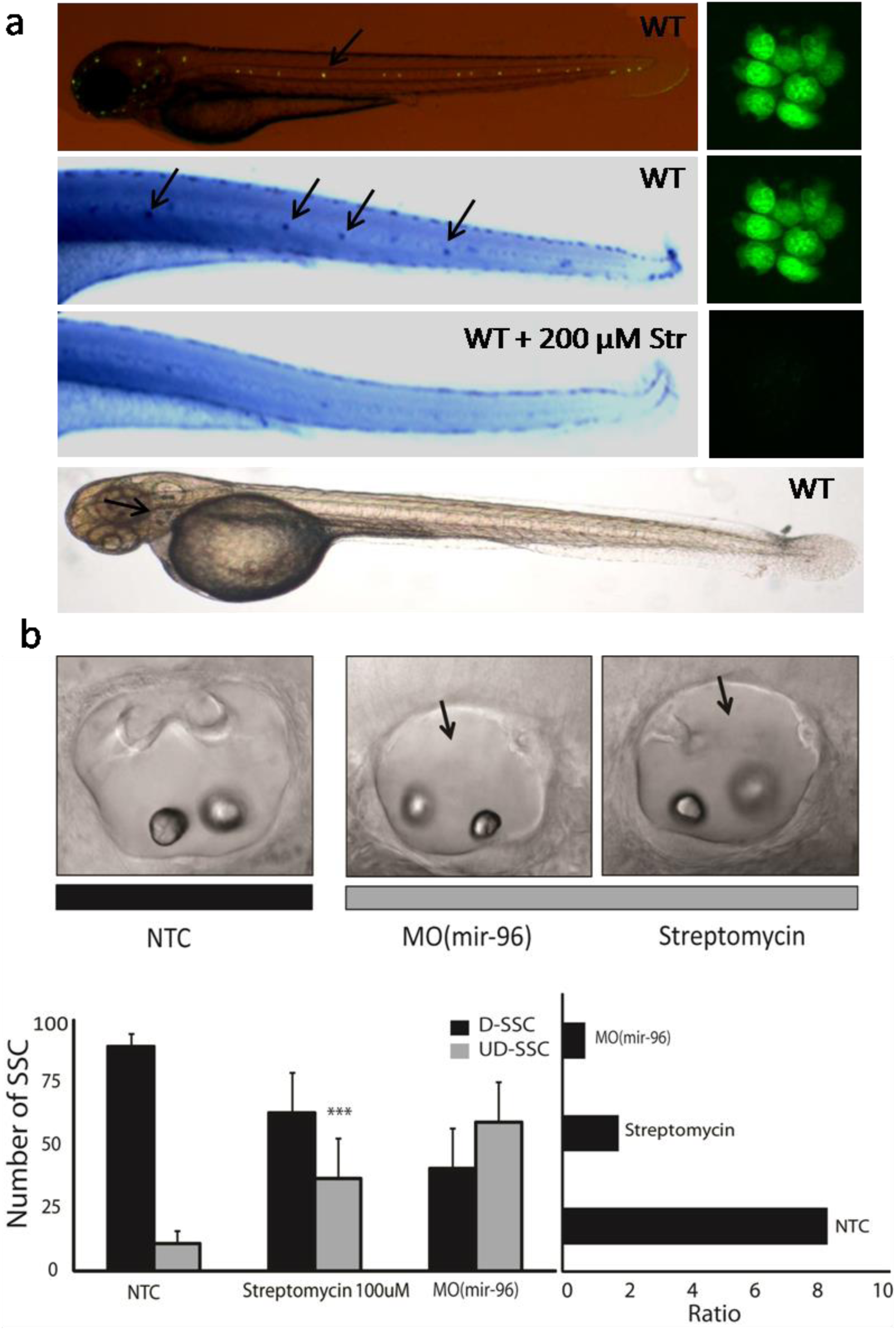
Hair cell bundle (Neuromast) status by DASPEI staining and whole mount *in-situ* hybridization of miR-96. a) Panel 1 -A fluorescence image overlaid with bright field of wildtypezebrafish embryos (5dpf) showing distribution of the hair cell bundles (neuromasts) along the lateral line. Side panel shows single neuromast stained with DASPEI. Panel 2 Whole mount *in-situ* hybridization of probes against miR-96 showing strong and specific expression in the neuromasts. Inset shows single neuromast stained with DASPEI.Panel 3 Upon treatment with 200μM, loss of signal of miR-96 is observed. Arrows indicate neuromasts of the lateral line.Panel 4- Indication of otolith location. b) Phenotypic profiling of zebrafish comparing Streptomycin treatment and miRNA knockdown. Upper panel - Representative bright field images showing differences in the otolith upon treatment. Streptomycin treatment at 100 μM mirrors the effect produced by double morphants (MO-96) created by knockdown of miR-96. Lower panel - The plot (left) shows observe a clear difference (*** - *P*<0.05, t-test) between the developmental pattern of the semicircular canals when animals are treated with the aminoglycosides when compared to the non-treated control sets, but similar to the morphants created by morpholino injection. The plot (left) are ratios (D/U) derived from the average of the absolute numbers for clarity of difference.

Further, we scored for similarities in developmental defects between morpholino based miRNA knockdown and Streptomycin treatment. It is well understood, that the anatomical features of mammalian and the zebrafish inner ears have a high degree of similarity(Whitfield, 2002). Particularly, the otic development programs and features of the otocysts have been earlier used to estimate the roles of specific miRNA in inner ear development. Otocyst formation is completed at the time window of 48hpf −52hpf(Haddon and Lewis, 1996) through the morphogenesis and formation of the otoliths and semicircular canals. In a normal developmental scenario, the superior semicircular canals (referred to SSC) (Fig 4a, last panel) develop at this stage and appear as craters in the otherwise smooth otocyst. Knockdown of miRNA from this cluster has been shown to deregulate this developmental process, giving a smooth appearance to the otocyst (Li, et al., 2010). We scored for the parallel effects of morpholino knockdown and aminoglycoside treatment. Morpholino based knockdown and aminoglycoside treatment at the single cell stage, both gave a drastic change in the ratio of fully developed and underdeveloped SSCs (Fig 4b). These results collectively indicate a profound phenotypic similarity between MO based knockdown and Streptomycin treatment in developing zebrafish embryos, suggesting that Streptomycin acts to decrease levels of this miR cluster and thus lead to observed hearing/balance defects.

Our data cumulatively suggest that ototoxicity of Streptomycin may funnel through physical interaction and subsequent reduction of miR-96 cluster levels. However, it is well documented that Streptomycin induced mitochondrial damage and subsequent apoptosis of hair cells explains the observed ototoxicity. This may occur due to the genetic predisposition of the patient due to certain mitochondrial mutation (Prezant, et al., 1993) among other factors. It is quite possible that our earlier results showing a signal loss of miR-96 in our transcriptional profiling and whole mount *in-situ* hybridization, may originate by complete loss of the hair cell as a whole due to mitochondrial damage and apoptotic cell death. To distinguish these effects, we rationalized that at low concentrations where hair cell viability (as a function of DASPEI based mitochondrial uptake) is preserved, if miR-96 and cluster miRNA levels are reduced, it would indicate a mitochondria-independent pathway of progressive ototoxicity of the drug. We thus, reduced the treatment concentrations progressively until, we reached an optimal concentration of 20μM where we did not observe an noticeable change in viability of hair cells (Fig 5a, b). *In-situ* hybridization showed signal of miRNA cluster members (Fig 5a, b). We then quantified miRNA levels with qRT-PCR at this concentration range. Indeed levels of miR- 96, miR-182 and miR-183 were reduced (Fig 5c).

**Figure 5.**
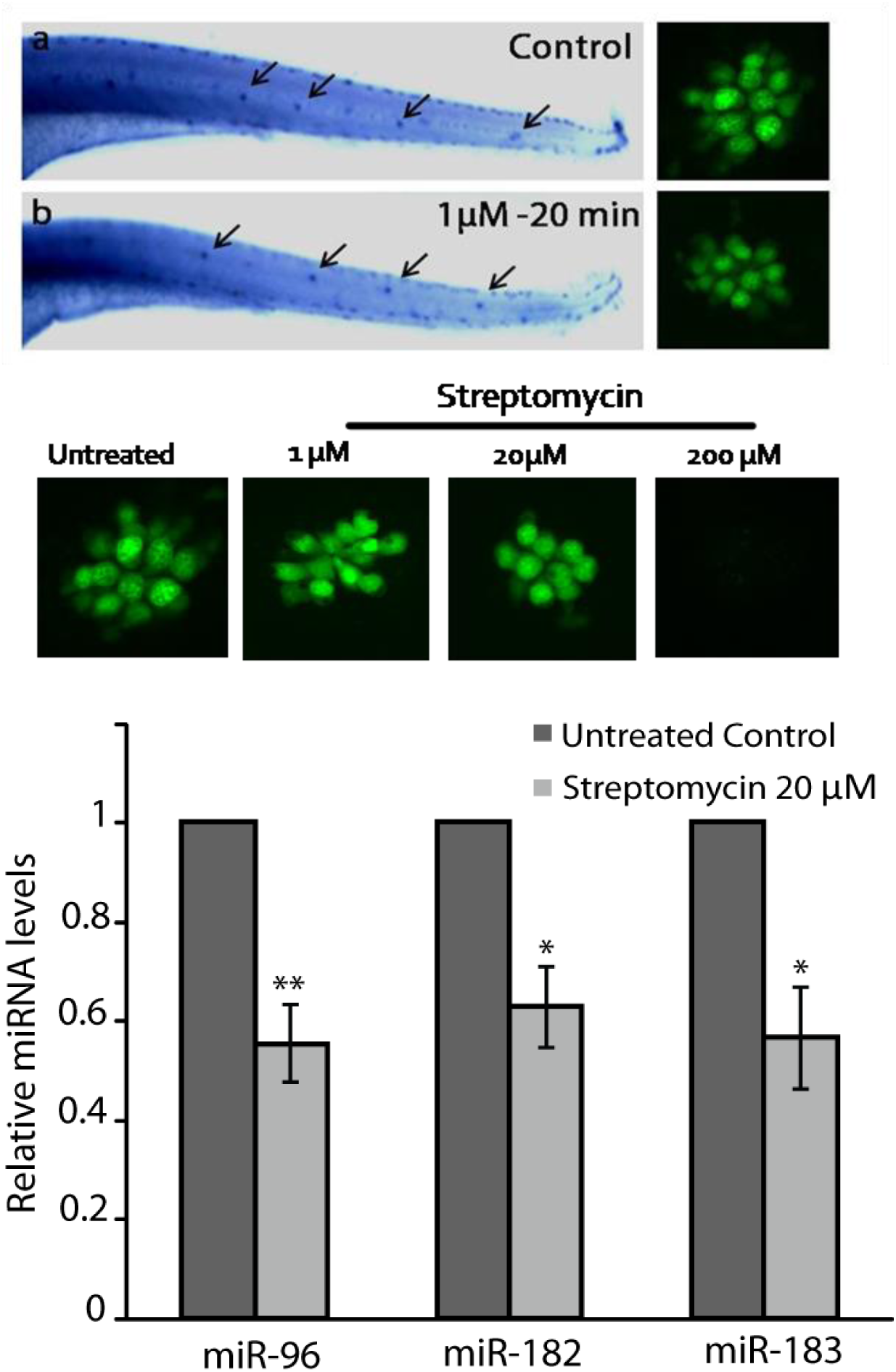
Hair cell bundle (neuromast) status reflecting mitochondrial health (DASPEI uptake) at lower optimal concentrations. a) Whole mount *in-situ* hybridization of probes to miR-96 showing localized expression to neuromasts. Right panel shows single neuromast stained with DASPEI. b) Subsequent treatment at a lower optimal concentration prevents signal loss from the neuromasts. Right panel shows comparable mitochondrial health to control non-treated mbryos. Increasing the concentration to 20μM does not decrease mitochondrial heath drastically (lower panel). c) qRT-PCR of miR-96 cluster members shows persistent decrease in levels of miRNA even at low optimal concentration of Streptomycin where mitochondrial health is preserved.

### Discussion

Drug toxicities until now were explained in the context of drugs causing side effects by mechanisms which generally involve mistargeting in the proteome. Several such examples have been documented in the past which have also given insight into the so called ‘polypharmacological’ potential to these drugs(Ashburn and Thor, 2004). Recent studies on ‘drug repositioning’ also have given opportunities to explore existing drug usage in the context of different diseases beyond what they were intended for. Aminoglycosides have already been used for promoting read through of premature termination codons in various conditions (Heier and DiDonato, 2009). Mirroring these efforts, and owing to the growing body of literature of druggable RNA we sought to explore the potential of well characterized aminoglycosides to modulate miRNA. Our results indicate that although aminoglycosides have a potential to modulate miRNA levels, their usage as direct candidates are limited owing to their miRnome wide perturbation. As indicated before, of the 6 aminoglycosides tested, Streptomycin, Gentamycin and Dihydrostreptomycin show a promising potential as starting scaffolds due to their relatively specific and minimal global perturbation. It is also evident, that not all perturbations may be a result of structural similarity calling for further intense scrutiny of potential secondary effects. Nevertheless, the extent of cross talk of a simple aminoglycoside antibiotic like Streptomycin is significant in the context of increasing ascription of function to the non-coding transcriptome which serves as a rich sink of secondary structures (Fig S8).

We further demonstrate this cross talk and the emergent explanation of drug toxicity of well characterized aminoglycoside antibiotics, funneling via perturbation of miRNA function. Using zebrafish as a model system, we show that ototoxicity of Streptomycin can channel by decreasing miR-96 and other cluster members. Here, we show that miR-96, miR-182 and miR- 183 (amongst other hearing related miRNA) decrease in expression due to Streptomycin administration. We also provide evidence that this decrease is due to the binding of the drug to the precursor miRNA, and subsequent perturbation of maturation by Dicer. Recently, a genetic study involving patients with non-syndromic inherited hearing loss showed that a novel mutation outside of the seed region of miR-96 (in the terminal loop) is strongly associated with the disease outcome. This mutation causes an altered stability of pre-miRNA causing a lowered maturation (Solda, et al., 2012) thus functionally causing the observed phenotype. This observation lends credible support to the results obtained in our study.

Although, mechanisms of ototoxicity of aminoglycosides have been previously described to stem from the interactions of aminoglycosides with various intracellular molecules like ferricions, phoshoinositides, mammalian mitochondrial RNA, and proteins (Karasawa and Steyger, 2011), it remains unclear to what degree to these individual components contribute to the toxicity observed. Recent evidences also point out that, even antibacterial effects of aminoglycosides can assume different trajectories (Foti, et al., 2012; Kohanski, et al., 2007; Kohanski, et al., 2008) involving various different pathways in a chronological manner, giving impetus to ask if aminoglycosides can perturb mammalian cell function in a similar fashion.

The severity of ototoxicity of Streptomycin differs with treatment dosages. Clinically, till date, only administration and dosage regimes have been modified without a clinically relevant remedial counter measure (Begg and Barclay, 1995; Mitnick, et al., 2008). Most of the interactions of aminoglycosides result in the generation of ROS and subsequent mitochondrial destabilization and apoptosis (Karasawa and Steyger, 2011). We here suggest, in the myriad complexities of overlapping pathways of ototoxicity, perturbation of miRNA function may be one of the important contributory factors, considering the fact that previous attempts of otoprotectant strategies (Huth, et al., 2011) did not completely ameliorate the ototoxicity of aminoglycosides. According to a recent report (Kalghatgi, et al., 2013) aminoglycosides induce mitochondrial dysfunction and damage. However in that study, ROS is claimed to be a contributor, but not the sole contributor of the resulting cellular damage, thus aligning well with the non-uniformity of the clinical success of anti-oxidants.

Although, the downstream effects of miRNA perturbation in this case is not clear yet, we suggest, that residual, local cytosolic concentrations of Streptomycin, may be enough to cause miRNA decrease and subsequently reduce efficacy of any otoprotectant. Our results thus indicate that although ROS induced damage may explain most cases of drug induced toxicity, it may not be the only determinant. This is evident from the fact that, all the drugs do not give the same expression change pattern (Fig 1a). In another example, the cardiotoxicity induced by administration of chemotherapy agent doxorubicin, was explained to be driven by a topoisomerase isoform *Top2B*(Zhang, et al., 2012), and not solely because of redox cycling. The primary killing mechanisms of bactericidal antibiotics involving ROS production were also put to debate (Keren, et al., 2013; Liu and Imlay, 2013), suggesting that the situation may not be so different for observed effects of aminoglycosides in eukaryotic systems. However follow up studies in prokaryotes (Belenky, et al., 2015; Dwyer, et al., 2014; Dwyer, et al., 2015) indicate that redox (ROS) related physiological alterations form a part of drug lethality, however it is not the sole contributor.

In summary this report suggests that RNA binding drugs need to be re-evaluated for the off targeting effects in backdrop of non-coding RNA. These results also imply that drug side effect evaluation should include a screen for such possibilities. Our results have exemplified that even classical antibiotics like Streptomycin have significant off targeting effects in the context of miRNA perturbation which can be used for potential drug repositioning upon further characterization and modifications. Our results also uncover a previously unanticipated mechanism of ototoxicity by an RNA binding drug which has been in use for over half a century.

## ACKNOWLEDGEMENT

The authors would like to express their gratitude to Dr.Koyeli Mapa, Dr.Soumya Sinha Roy and other members of the Maiti lab for their insightful discussions and comments on the manuscript. The authors acknowledge Dr. Satyaprakash Pandey for his help with the footprinting experiments. GGJ and SR acknowledge CSIR for fellowship.

### FUNDING

This work was supported by the Council of Scientific and Industrial Research (CSIR), Government of India - network project BSC0123 and the Swarnajayanti Fellowship, Government of India to SM. Funding for open access charge: Council of Scientific and Industrial Research (CSIR).

## EXPERIMENTAL PROCEDURES

### Cell culture, miRNome expression profiling, qRT-PCR

The MCF-7 cells were passaged in 24-well plate (5×10^4^ cells/well), grown to 70% con?uency and treated with different aminoglycosides at a concentration of 10 μM. Cells were then incubated at 37 °C for 24 h (DMEM, 5% CO_2_). The media was removed, and cells were washed with PBS buffer and total RNA was isolated using Trizol^®^ Reagent (Invitrogen). Expressions of mature microRNAs (380, plate 1) were determined by Human miRNome profiler kit. (SBI, catalog number RA660A-1). Primer sequences are available on request. The microRNA expression levels were detected using SYBR- fast 2X mastermix (KAPA Biosystems, KM4101) on Roche Lightcycler LC 480. All experiments were repeated twice and Ct values differing beyond a range of ± 0.5 were discarded. All the data were then analyzed using the ΔΔ Ct method (Schmittgen and Livak, 2008). For the RNA samples isolated from zebrafish embryos, an in house custom array in a 384 well format was created in the same format, allowing global quantification of miRnome (248 miRNA, mirbase V15) in D.rerio.

### Undirected Network Construction

For ease of interpretation and visualization of the transcriptional response data, the dataset after applying a cutoff of 2 fold for downregulation was converted into an adjacency matrix. The interactions of each small molecule drug with given miRNA were computed using a custom in house script. The network was then visualized using Gephi (version 0.8.2 beta).

### Footprinting and Dicer Cleavage Assay

P^32^-5′end labeled in-vitro transcribed pre-miRNA was subjected to a denaturation and renaturation procedure before structure probing. The P^32^-5′end-labeled RNA (60000 cpm) was incubated in 50 mMTris-HCl (pH 7.5), 50 mMNaCl, 5mM MgCl2 in a total volume of 8 μL. The sample was heated at 90°C for 1 min and left in a thermal block for slow (∼10 min) cooling to 37°C. 1 μL Streptomycin was added to a final concentration of 10 μM and incubated at 37°C for 10 minutes. For S1 nuclease probing 1μl of S1 nuclease was added (total amount of enzyme is each reaction is 47.5 U) and incubated at 37°C for 1 min. The reaction was stopped by adding 10 μL of 95% formamide with dyes. 10 μL was loaded in 15% denaturing gel. For T1 nuclease probing 1uL of T1 nuclease was added (total amount of enzyme is each reaction is 0.05 U) and incubated at 37°C for 1.5 minute. The reaction was stopped by adding 10 μL of 95% formamide with dyes. Alkaline digestion ladder and RNase T1 digestion ladder was generated as described earlier. Samples were run in 15% denaturing PAGE and exposed to phosphor screen overnight. Cleavage profiles were generated using Multi Gauge software (version 2.3).

The Dicer cleavage assay was performed at 37 °C for 4 h using Dicer buffer (Invitrogen). Each reaction mixture contained 3 μL of P^32^-5′end labeled pre-miR (∼20 ng), and 0.01 U of Dicer (Invitrogen). We have used 10μM aminoglycoside in a final volume of 10 μL. The reaction was stopped by Dicer stop buffer (Invitrogen). Equal volume of 95% formamide with dyes was added, loaded in 15% denaturing gel.

### Surface Plasmon Resonance

All kinetic parameters of binding of the peptides to RNA were calculated from SPR experiments performed on BIAcore 3000 system running with BIAcore 3000 control software version 4.1.2. The pre-miRNA were biotinylated at the 5′ end of the RNA. The 100 nM stock solution of RNA was used for immobilization on the flow cell 2 of streptavidin coated SA chip until an RU change of 500-600 was achieved. After immobilization the buffer was allowed to flow on the sensor chip surface to remove any unbound RNA. Flow cell 1 was left blank to account for nonspecific background signal, and was subtracted from the signal in flow cell 2. All the RNA Streptomycin solutions were prepared in 10 mM HEPES (pH 7.5) buffer with 10 mM NaCl and 1 mM MgCl_2_. Serial dilutions of the 10 μM stock were done to make their concentration series. The binding and dissociation were monitored for 300 s, followed by 60 s regeneration using buffer. The sensograms were analysed with BIAevaluation software version 4.1.1 using 1:1 Langmuir model and the goodness of the fitting was monitored by χ2 value.

### Zebrafish Husbandry and Treatment with Aminoglycosides

Zebrafish used in this study were housed at the CSIR-Institute of Genomics and Integrative Biology following standard husbandry practices. Zebrafish embryos were obtained by pair- wise mating of adult fish. Zebrafish experiments were performed in strict accordance with the recommendations and guidelines laid down by the CSIR-Institute of Genomics and Integrative Biology, India. The protocol was approved by the Institutional Animal Ethics Committee (IAEC) of the CSIR Institute of Genomics and Integrative Biology, India (Proposal No 45a). Aminoglycosides were diluted in defined E2 embryo medium. Animals were treated with drug or embryo media (mock-treated controls) for times indicated, subsequently washed rapidly three times in fresh embryo medium and allowed to recover for one hour.

### Microinjections

Morpholino injections were performed following a published protocols. Glass capillary (World Precision) micropipettes were pulled using Sutter Instrument (USA) and clipped appropriately for microinjection into one cell stage zebrafish embryos. Morpholinos (MO) were administered as 2-3 nanoliter (nl) injections containing 10 mM MO was injected at one cell stage zebrafish embryos.

### Whole mount in-situ hybridization and imaging

Whole-mount in-situ hybridiza-tion (WMISH) was performed as described previously (Kloosterman et al., 2006) 44. Embryos older than 22 hpf were treated with phenylthiourea (0.2mM) to inhibit pigmentation. Embryos were fixed in 4% paraformaldehyde in PBS at 4°C overnight or at room temperature for 2 h, followed by dehydration in methanol at 20°C for at least 2 h. After rehydration through graded methanols and treatment with proteinase K (2.8–22μg/ml), embryos were preincubated in hybridization buffer for 2–3 h and then were hybridized with antisense RNA probes (1 μg /ml) at 65°C or lockednucleic acid (LNA)- modified DNA probes for miRNAs (20 nM; Exiqon) at 51°C overnight. The 3′ ends of LNA-modified probes were labeled with digoxygenin (DIG; Roche Applied Science) according to the manufacturer’s recommendation before hybridization. After hybridization and high-stringency washes, embryos were blocked in 2% goat serum and 2 mg/ml BSA in 0.1% Tween 20 in PBS at roomtemperature for 1 h and incubated in anti-DIG conjugated to alkaline phosphatase antibody (Roche Applied Science) at 4°Covernight. Embryos were incubated in BCIP (5-bromo-4-chloro-3′-indolyl phosphate p-toluidine salt; 175μg/ml) and NBT (nitro- blue tetrazolium chloride; 225 μg/ml) (Roche Applied Science) at room temperature for2h or at 4°C overnight for purple colour reaction.

### Hair cell imaging

Post dye staining, whole animal embryos were imaged using an upright Zeiss Axioscope 40 microscope (Carl Zeiss, Germany) using 2.5X and 5X magnification and 0.075 numerical aperture. To visualize the hair cell bundles specifically, confocal microscopy using a Zeiss LSM 510 Meta system at 40X magnification with the appropriate filter sets and uniform settings for all animals imaged per set. All images were processed using Adobe software.

---

**Table 1.** Table summarizes binding of aminoglycosides to the pre-miRNA which were synthesized by in vitro transcription as per manufacturer’s instructions (Ambion.,Inc). The summary is a result of two independent observations for each aminoglycoside to determine their physical binding to the respective precursor miRNA. Here only Streptomycin showed significant protection and thus binding to pre-mir-182 and pre-mir-96.

## REFERENCES

Ashburn, T.T., and Thor, K.B. (2004). Drug repositioning: identifying and developing new uses for existing drugs. Nat Rev Drug Discov 3, 673–683.

Bartel, D.P. (2009). MicroRNAs: target recognition and regulatory functions. Cell 136, 215–233.

Begg, E.J., and Barclay, M.L. (1995). Aminoglycosides-50 years on. Br J Clin Pharmacol 39, 597–603.

Belenky, P., Ye, J.D., Porter, C.B., Cohen, N.R., Lobritz, M.A., Ferrante, T., Jain, S., Korry, B.J., Schwarz, E.G., Walker, G.C., et al. (2015). Bactericidal Antibiotics Induce Toxic Metabolic Perturbations that Lead to Cellular Damage. Cell Rep 13, 968–980.

Bose, D., Jayaraj, G., Suryawanshi, H., Agarwala, P., Pore, S.K., Banerjee, R., and Maiti, S. (2012). The tuberculosis drug streptomycin as a potential cancer therapeutic: inhibition of miR-21 function by directly targeting its precursor. Angew Chem Int Ed Engl 51, 1019–1023.

Bowman, T.V., and Zon, L.I. (2010). Swimming into the future of drug discovery: in vivo chemical screens in zebrafish. ACS Chem Biol 5, 159–161.

Dwyer, D.J., Belenky, P.A., Yang, J.H., MacDonald, I.C., Martell, J.D., Takahashi, N., Chan, C.T., Lobritz, M.A., Braff, D., Schwarz, E.G., et al. (2014). Antibiotics induce redox-related physiological alterations as part of their lethality. Proc Natl Acad Sci U S A 111, E2100–2109.

Dwyer, D.J., Collins, J.J., and Walker, G.C. (2015). Unraveling the physiological complexities of antibiotic lethality. Annu Rev Pharmacol Toxicol 55, 313–332.

Emde, A., and Hornstein, E. (2014). miRNAs at the interface of cellular stress and disease. EMBO J 33, 1428–1437.

Foti, J.J., Devadoss, B., Winkler, J.A., Collins, J.J., and Walker, G.C. (2012). Oxidation of the guanine nucleotide pool underlies cell death by bactericidal antibiotics. Science 336, 315–319.

Francis, S.P., Katz, J., Fanning, K.D., Harris, K.A., Nicholas, B.D., Lacy, M., Pagana, J., Agris, P.F., and Shin, J.B. (2013). A novel role of cytosolic protein synthesis inhibition in aminoglycoside ototoxicity. J Neurosci 33, 3079–3093.

Guttilla, I.K., and White, B.A. (2009). Coordinate regulation of FOXO1 by miR-27a, miR- 96, and miR-182 in breast cancer cellsx. J Biol Chem 284, 23204–23216.

Haddon, C., and Lewis, J. (1996). Early ear development in the embryo of the zebrafish, Danio rerio. J Comp Neurol 365, 113–128.

Heier, C.R., and DiDonato, C.J. (2009). Translational readthrough by the aminoglycoside geneticin (G418) modulates SMN stability in vitro and improves motor function in SMA mice in vivo. Hum Mol Genet 18, 1310–1322.

Hobbie, S.N., Akshay, S., Kalapala, S.K., Bruell, C.M., Shcherbakov, D., and Bottger, E.C. (2008). Genetic analysis of interactions with eukaryotic rRNA identify the mitoribosome as target in aminoglycoside ototoxicity. Proc Natl Acad Sci U S A 105, 20888–20893.

Hong, W., Zeng, J., and Xie, J. (2014). Antibiotic drugs targeting bacterial RNAs. Acta Pharm Sin B 4, 258–265.

Horibe, T., Matsui, H., Tanaka, M., Nagai, H., Yamaguchi, Y., Kato, K., and Kikuchi, M. (2004). Gentamicin binds to the lectin site of calreticulin and inhibits its chaperone activity. Biochem Biophys Res Commun 323, 281–287.

Huth, M.E., Ricci, A.J., and Cheng, A.G. (2011). Mechanisms of aminoglycoside ototoxicity and targets of hair cell protection. Int J Otolaryngol 2011, 937861.

Jacquier, A. (2009). The complex eukaryotic transcriptome: unexpected pervasive transcription and novel small RNAs. Nat Rev Genet 10, 833–844.

Jayaraj, G.G., Nahar, S., and Maiti, S. (2015). Nonconventional chemical inhibitors of microRNA: therapeutic scope. Chem Commun (Camb) 51, 820–831.

Kalghatgi, S., Spina, C.S., Costello, J.C., Liesa, M., Morones-Ramirez, J.R., Slomovic, S., Molina, A., Shirihai, O.S., and Collins, J.J. (2013). Bactericidal antibiotics induce mitochondrial dysfunction and oxidative damage in Mammalian cells. Sci Transl Med 5, 192ra185.

Karasawa, T., and Steyger, P.S. (2011). Intracellular mechanisms of aminoglycoside-induced cytotoxicity. Integr Biol (Camb) 3, 879–886.

Karasawa, T., Wang, Q., David, L.L., and Steyger, P.S. (2011). Calreticulin binds to gentamicin and reduces drug-induced ototoxicity. Toxicol Sci 124, 378–387.

Karasawa, T., Wang, Q., David, L.L., and Steyger, P.S. (2010). CLIMP-63 is a gentamicin- binding protein that is involved in drug-induced cytotoxicity. Cell Death Dis 1, e102.

Keren, I., Wu, Y., Inocencio, J., Mulcahy, L.R., and Lewis, K. (2013). Killing by bactericidal antibiotics does not depend on reactive oxygen species. Science 339, 1213–1216.

Kohanski, M.A., Dwyer, D.J., Hayete, B., Lawrence, C.A., and Collins, J.J. (2007). A common mechanism of cellular death induced by bactericidal antibiotics. Cell 130, 797–810.

Kohanski, M.A., Dwyer, D.J., Wierzbowski, J., Cottarel, G., and Collins, J.J. (2008). Mistranslation of membrane proteins and two-component system activation trigger antibiotic-mediated cell death. Cell 135, 679–690.

Krol, J., Loedige, I., and Filipowicz, W. (2010). The widespread regulation of microRNA biogenesis, function and decay. Nat Rev Genet 11, 597–610.

Leclerc, D., Melancon, P., and Brakier-Gingras, L. (1991). Mutations in the 915 region of Escherichia coli 16S ribosomal RNA reduce the binding of streptomycin to the ribosome. Nucleic Acids Res 19, 3973–3977.

Leung, A.K., and Sharp, P.A. (2010). MicroRNA functions in stress responses. Mol Cell 40, 205–215.

Lewis, M.A., Quint, E., Glazier, A.M., Fuchs, H., De Angelis, M.H., Langford, C., van Dongen, S., Abreu-Goodger, C., Piipari, M., Redshaw, N., et al. (2009). An ENU-induced mutation of miR-96 associated with progressive hearing loss in mice. Nat Genet 41, 614–618.

Li, H., Kloosterman, W., and Fekete, D.M. (2010). MicroRNA-183 family members regulate sensorineural fates in the inner ear. J Neurosci 30, 3254–3263.

Liu, Y., and Imlay, J.A. (2013). Cell death from antibiotics without the involvement of reactive oxygen species. Science 339, 1210–1213.

Mencia, A., Modamio-Hoybjor, S., Redshaw, N., Morin, M., Mayo-Merino, F., Olavarrieta, L., Aguirre, L.A., del Castillo, I., Steel, K.P., Dalmay, T., et al. (2009). Mutations in the seed region of human miR-96 are responsible for nonsyndromic progressive hearing loss. Nat Genet 41, 609–613.

Mendell, J.T., and Olson, E.N. (2012). MicroRNAs in stress signaling and human disease. Cell 148, 1172–1187.

Mitnick, C.D., Shin, S.S., Seung, K.J., Rich, M.L., Atwood, S.S., Furin, J.J., Fitzmaurice, G.M., Alcantara Viru, F.A., Appleton, S.C., Bayona, J.N., et al. (2008). Comprehensive treatment of extensively drug-resistant tuberculosis. N Engl J Med 359, 563–574.

Miyazaki, T., Sagawa, R., Honma, T., Noguchi, S., Harada, T., Komatsuda, A., Ohtani, H., Wakui, H., Sawada, K., Otaka, M., et al. (2004). 73-kDa molecular chaperone HSP73 is a direct target of antibiotic gentamicin. J Biol Chem 279, 17295–17300.

Oishi, N., Duscha, S., Boukari, H., Meyer, M., Xie, J., Wei, G., Schrepfer, T., Roschitzki, B., Boettger, E.C., and Schacht, J. (2015). XBP1 mitigates aminoglycoside-induced endoplasmic reticulum stress and neuronal cell death. Cell Death Dis 6, e1763.

Owens, K.N., Santos, F., Roberts, B., Linbo, T., Coffin, A.B., Knisely, A.J., Simon, J.A., Rubel, E.W., and Raible, D.W. (2008). Identification of genetic and chemical modulators of zebrafish mechanosensory hair cell death. PLoS Genet 4, e1000020.

Prezant, T.R., Agapian, J.V., Bohlman, M.C., Bu, X.D., Oztas, S., Qiu, W.Q., Arnos, K.S., Cortopassi, G.A., Jaber, L., Rotter, J.I., et al. (1993). Mitochondrial Ribosomal-Rna Mutation Associated with Both Antibiotic-Induced and Non-Syndromic Deafness. Nature Genetics 4, 289–294.

Rudnicki, A., and Avraham, K.B. (2012). microRNAs: the art of silencing in the ear. EMBO Mol Med 4, 849–859.

Schmittgen, T.D., and Livak, K.J. (2008). Analyzing real-time PCR data by the comparative C(T) method. Nat Protoc 3, 1101–1108.

Shulman, E., Belakhov, V., Wei, G., Kendall, A., Meyron-Holtz, E.G., Ben-Shachar, D., Schacht, J., and Baasov, T. (2014). Designer aminoglycosides that selectively inhibit cytoplasmic rather than mitochondrial ribosomes show decreased ototoxicity: a strategy for the treatment of genetic diseases. J Biol Chem 289, 2318–2330.

Solda, G., Robusto, M., Primignani, P., Castorina, P., Benzoni, E., Cesarani, A., Ambrosetti, U., Asselta, R., and Duga, S. (2012). A novel mutation within the MIR96 gene causes non-syndromic inherited hearing loss in an Italian family by altering pre-miRNA processing. Hum Mol Genet 21, 577–585.

Sood, P., Krek, A., Zavolan, M., Macino, G., and Rajewsky, N. (2006). Cell-type-specific signatures of microRNAs on target mRNA expression. Proc Natl Acad Sci U S A 103, 2746–2751.

Ushakov, K., Rudnicki, A., and Avraham, K.B. (2013). MicroRNAs in sensorineural diseases of the ear. Front Mol Neurosci 6, 52.

Whitfield, T.T. (2002). Zebrafish as a model for hearing and deafness. J Neurobiol 53, 157–171.

Will, S., Joshi, T., Hofacker, I.L., Stadler, P.F., and Backofen, R. (2012). LocARNA-P: accurate boundary prediction and improved detection of structural RNAs. RNA 18, 900–914.

Zhang, S., Liu, X., Bawa-Khalfe, T., Lu, L.S., Lyu, Y.L., Liu, L.F., and Yeh, E.T. (2012). Identification of the molecular basis of doxorubicin-induced cardiotoxicity. Nat Med 18, 1639–1642.

